# Gravitropic Movement in Wheat Coleoptile is Regulated by Ultradian Growth Oscillations

**DOI:** 10.1101/093146

**Authors:** Renaud Bastien, Olivia Guayasamin, Stéphane Douady, Bruno Moulia

## Abstract

To stand straight and upright along their growth, plants needs to regulate actively their posture. Gravitropic movement, which occurs when plants modify their growth and curvature to orient their aerial organ against the force of gravity, is a major feature of this postural control. A recent model has shown that graviception and proprioception are sufficient to account for the gravitropic movement and subsequent organ posture demonstrated by a range of species. However, some plants, including wheat coleoptiles, exhibit a stronger regulation of posture than predicted by the model. Here, we performed an extensive kinematics study on wheat coleoptiles during a gravitropic perturbation experiment in order to better understand this unexpectedly strong regulation. Close temporal observation of the data revealed that both perturbed and unperturbed coleoptiles showed oscillatory pulses of elongation and curvature variation that propagated from the apex to the base of their aerial organs. In perturbed (tilted) coleoptiles, we discovered a non-trivial coupling between the oscillatory dynamics of curvature and elongation. This relationship appears to be critical to the postural control of the organ, and indicates the presence of a mechanism that is capable of affecting the relationship between elongation rate, differential growth, and curvature.

## Introduction

Plants are living organisms that have long been observed to adaptively regulate their shape and movement in order to optimize their physical performance. The best-known example of an adaptive plant response is phototropism, which describes the ability of plants to track sunlight in order to maximize photosynthesis. While phototropism often results in plants growing upwards toward the sun, it is not the only mechanism by which plants can achieve upward orientation and growth. In fact, many plants exhibit gravitropism and can spatially orient their organs with the gravitational field [1, 2], yielding roots that grow downward in direction of gravity and stems elongating upwards counter to gravitational pull. Gravitropism, phototropism, and other forms of plant tropism result from an asymmetrical distribution of auxin in the elongation zone of various organs [2–4]. The unequal distribution of auxin in these regions causes a difference in growth rate between opposite sides of the plant organ, resulting in a curvature (Figure 1) that orients the plant organ toward a salient stimulus [3, 5–7].

**Fig 1.**
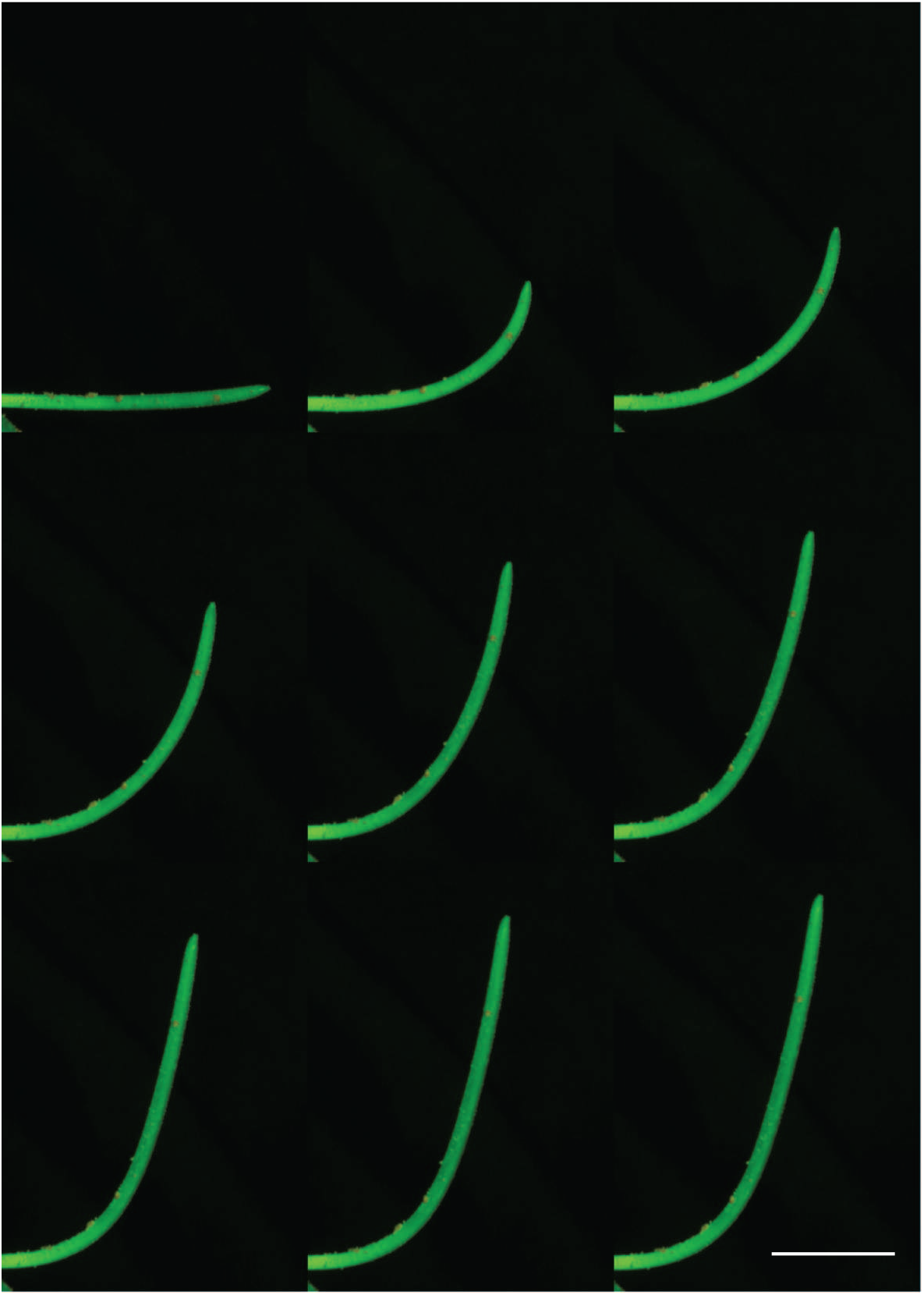
Straightening movement of a wheat coleoptile, in the standard reading direction. The white line accounts for 10 mm. The time between each frame is 150 minutes. The orange points on the coleoptile are the fluorescent markers used for the measurement of growth. Movies of all plants are available online [8, 9].

Studies on the kinematics of gravitropic movement in plants have largely neglected the aerial organs, instead focusing on roots as they are less affected by phototropism [10–12]. In roots, graviperception occurs only in the root cap and this signal should be transmitted to the underlying elongation zone. Subsequently the regulation of the movement is limited to the apical part [13]. Furthermore, unlike aerial organs, roots rely on physical contact with the surrounding substrate in order to help regulate their posture.

In contrast to roots, both perception and the control of the movement are possible along the entire length of the aerial stem [5, 13, 14]. Because they are not embedded within a substrate, these freestanding aerial organs require internal mechanisms to regulate their posture and spatial orientation in the absence of light. In order to consistently grow straight upwards, aerial organs must exhibit both graviception and proprioception, the ability of plants to perceive their own shape and respond accordingly [2, 14, 15]. These two perceptions have different effects on plant movement and shape: graviception tends to increase the curvature of the organ, allowing it to reach vertical while proprioception causes plants to modify their growth in order to remain straight [14]. Aerial organs must, therefore, balance the effects of graviception and proprioception in order to successfully grow upwards against the force of gravity. For these reasons, studying gravitropism in plant aerial organs provides a unique opportunity to understand the how plants integrate multi-sensory information and regulate growth and movement.

Recently, the process of gravitropic movement in plant aerial organs has been formalized in a unifying model called the “AC model” [5, 14], where the additive (but opposing) effects of graviception and proprioception are expressed by

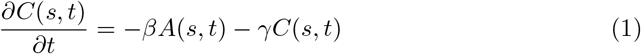

where *s* is the position along the organ, from the apex to the base, *t* is the time, *A*(*s, t*) is the local angle of the organ to the vertical, and *C*(*s, t*) is the local curvature (i.e. the rate of change of *A*(*s, t*) along *s, ∂_s_A*(*s, t*) = *C*(*s, t*)). The parameters *β* and *γ* respectively represents the organ’s graviceptive and proprioceptive sensitivities. In this model, the dynamics of aerial stem movement are represented by a unique dimensionless number, *B* the “balance number” described by

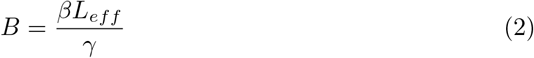

where *L_eff_* is the effective length of the organ. If the length of the organ *L* is greater than the length of the growth zone (*L_gz_ > L*) at the time before stimulation *L*(*t* = 0), then *L_eff_* = *L*. In contrast, if (*L_gz_ < L*) at *L*(*t* = 0), then *L_eff_* = *L_gz_* (see Figure 2 for a full visual depiction). The number *B* expreses the ratio between graviception, *β*, and proprioception γ, scaled to the length of the organ that can move. Because of this, the AC model is also known as the “gravipropriceptive” model. If *B* is small, then proprioception tends to dominate the movement and the organ does not reach the vertical, while if *B* is large the organ oscillates around the vertical until the apical tip of the aerial organ is aligned with the gravity field. Because *B* is a dimensionless value that is readily quantifiable from simple morphometric experiments, it can be used to make quantitative comparisons across a variety of model systems. Indeed, this model is able to explain the complex kinematics of gravitropism in at least eleven species covering a broad taxonomic range of angiosperms, habitats, and aerial organ types, suggesting that it captures a fundamental regulatory process of plant tropism [5, 14]. The *AC* model has been extended to separate the *AC* regulatory scheme from the growth activity powering the tropic movement, giving rise to the *ACE* model [5]. This study revealed that postural control in actively elongating organs is a challenge. But all the studied species were found to have adapted to it within the *AC* regulatory scheme, though sufficient proprioceptive sensitivity.

**Fig 2.**
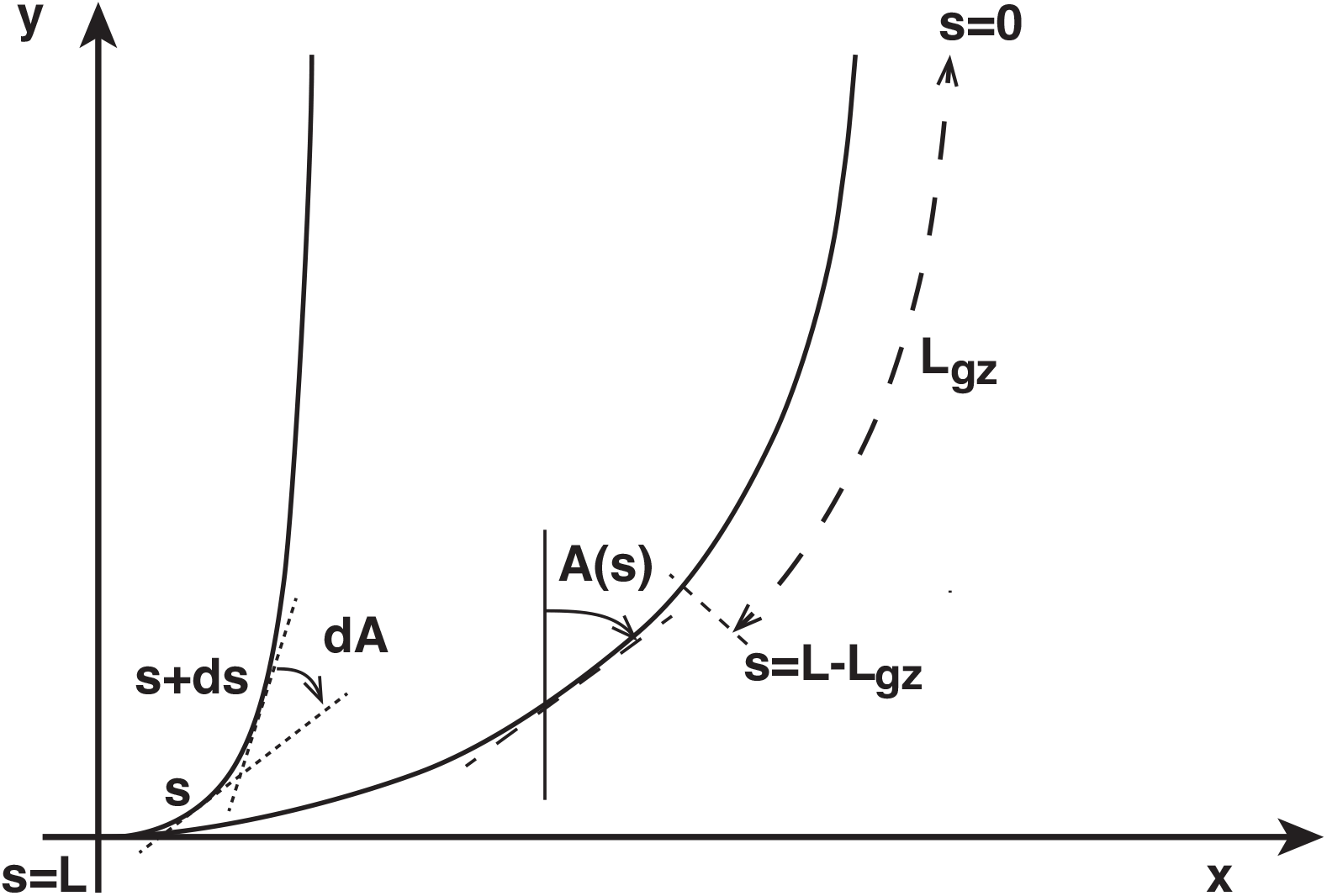
Geometric description of the organ. We define a set of coordinates (*x,y*) in the plane that intersects with the organ. The vertical direction *y* is aligned with the gravity field. The median line of an organ of total length L is defined by its curvilinear abscissa *s*, with *s* = 0 referring to the apex and *s* = *L*(*t*) referring to the base. In an elongating organ, only the part inside the growth zone, of length *L_gz_* from the apex, is able to curve (with *L_gz_* = *L* at early stages and *L_gz_* < *L* later on [5]). The local orientation of the organ *A*(*s, t*) is defined at each point along the median with respect to the vertical. Two curves are represented here with the same apical angle *A*(0, *t*) = 0 but different shapes. The measurement of *A*(*s, t*) along the entire median line is necessary to specify the full shape. Due to the symmetry of the system around the vertical axis, the angle *A*(*s, t*) is a zenith angle, zero when the organ is vertical and upright. Clockwise angles are considered positive.

However, many plants seem to present a more comprehensive regulation of gravitropic movement than currently described by the AC model. In particular, this model often predicts that aerial organs will “overshoot” the vertical (Figure 3), which occurs when the curvature of the aerial organ is too great and apical tip crosses over the vertical. In contrast to these predictions, many plants do not express a physical overshoot either during movement or in their final shape [14, 16], indicating that they possess an additional regulatory mechanism currently unrepresented in the AC model. One reason for the failure to include this term in the AC model may be that this regulatory mechanism can not be properly observed without clear and complete observations of plant movement, and all-inclusive plant kinematics are often not quantified or reported. Measurements of plant movement that focus on the first instants of the gravitropic reaction [17–19], or restrict analysis to the movement of the apical tip alone [18–20], give an incomplete picture of the kinematics of gravitropic movement and are insufficient for understanding its regulation [3, 13, 14, 21].

To discover the additional movement regulations that prevent plants from overshooting the vertical, we recorded the complete kinematics of wheat *(Triticum aestivum)* coleoptiles during a gravitropic experimental study. Wheat coleoptiles were chosen as our model system because they can be grown in absence of light, eliminating the concern of phototropic effects. Additionally, they have not been observed to overshoot the vertical [5] making them likely candidates for detecting a mechanism that prevents overshoot. After allowing seeds of wheat to germinate in a vertical position until they matured into coleoptiles, individual plants were subjected to one of two spatial reorientation conditions: no spatial reorientation (vertical) or 90^°^ reorientation (horizontal).Their movements were then observed for 24 hours.

**Fig 3.**
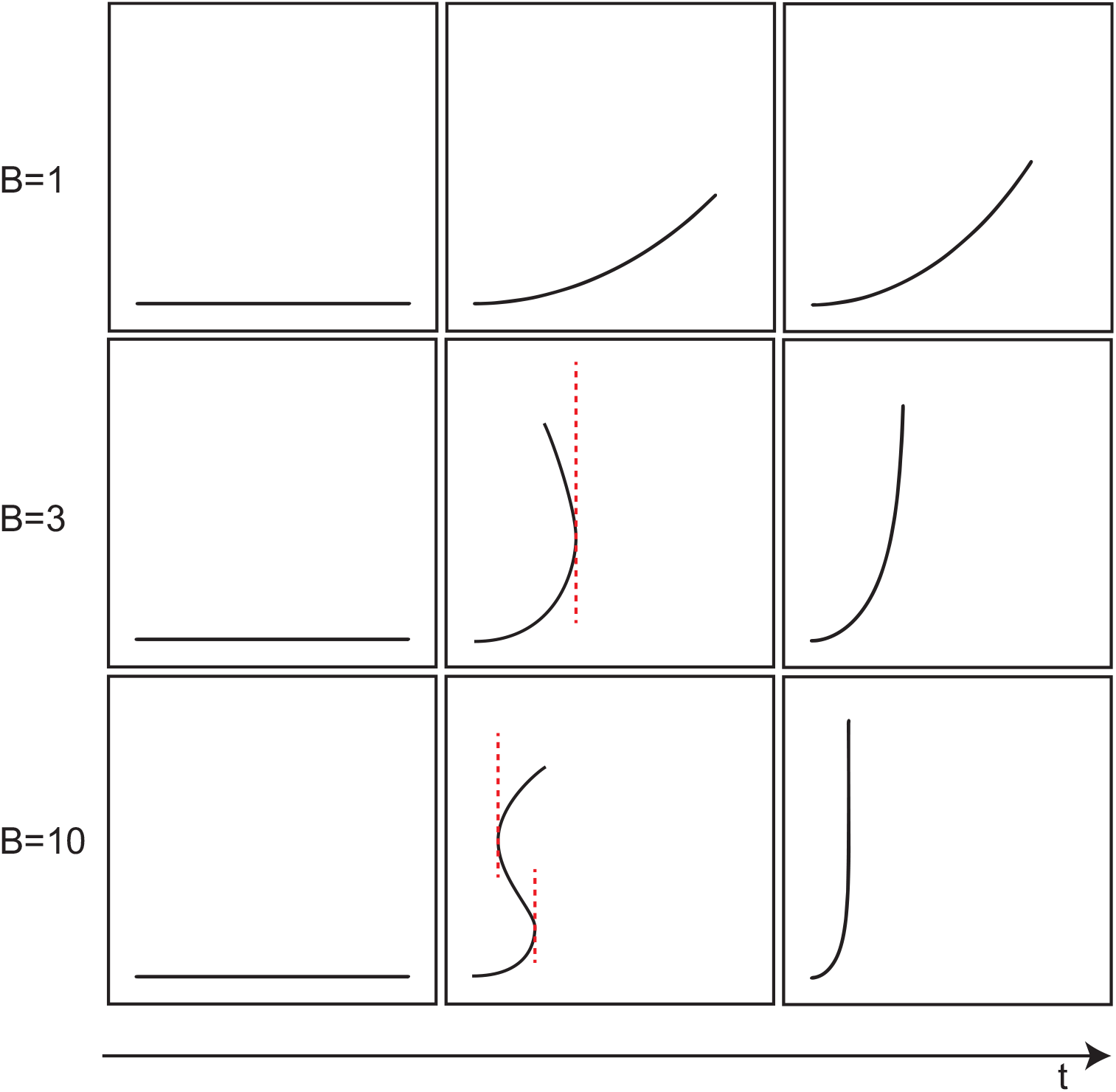
Straightening movements of a tilted virtual plants as observed in the *AC* model. From top to bottom, the values of B are increasing (1, 3, 10). From left to right, initial state, the virtual plant is straight and tilted 90° from the vertical, an intermediary state and finally, the equilibrium state where no more movements observed. As the value of B is increased, overshoots of the vertical are observed in the intermediary state (*B* = 3 C-shape, *B* = 10 S-shape). The length of the curve of the zone is getting smaller as the value of *B* is increased.

Throughout the observation period we recorded and quantified the growth and structure of the entire coleoptile with KymoRod, a new tool developed to extract the complete kinematics of an aerial organ in movement [22]. Using this approach, we compared the kinematics of disturbed and undisturbed coleoptiles in order to discover the mechanism responsible for the regulation of gravitropic movement beyond what is included in the AC model. As these dynamics are not yet described or defined in the current formulation of the AC model, we further seek to do so here.

## Materials and Methods

### Coleoptile Preparation and General Methods

Seeds of recital wheat (*Triticum aestivum)* were immersed in water for 6 hours to initiate germination, then each seed was attached to a plastic test tube filled with cotton and water. Because wheat coleoptiles do not sprout vertically from the seed’s longitudinal axis, all seeds were adhered at a 45° angle to ensure that coleoptiles would grow vertically with minimal initial curvature. After being attached to test tubes, all seeds were put in a darkroom to germinate for 3 to 4 days until coleoptiles were 10 — 20mm long. Each coleoptile was then dusted with an orange, non-toxic, fluorescent dry painting pigment *(Sennelier 648)*, leaving “tracer” particles on the coleoptile surface that were later used for extracting kinematics (Figure 4).

**Fig 4.**
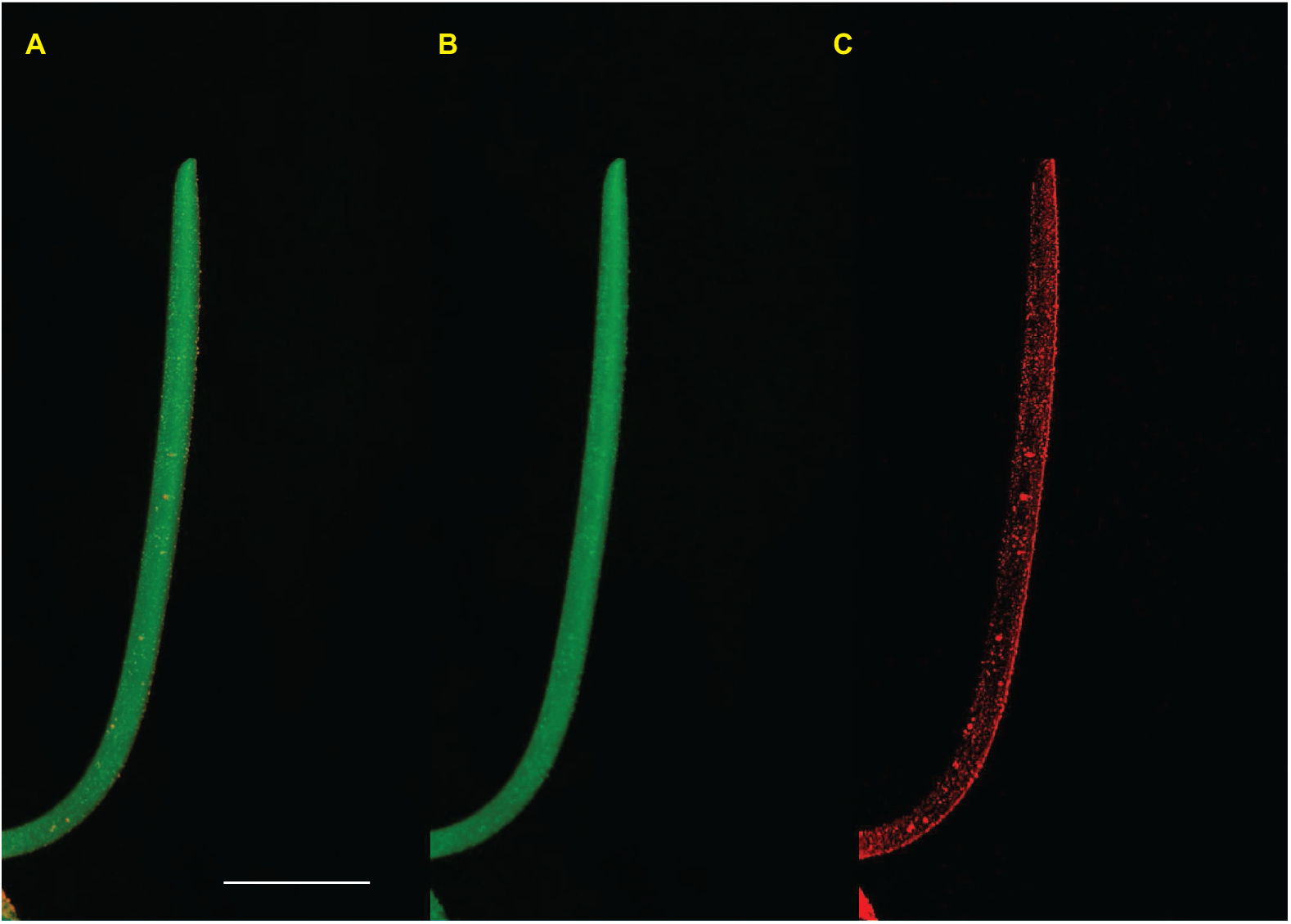
Picture of a coleoptile obtained from experiments. The white line is 1cm. Pictures are taken in the absence of light with the flash light of the camera filtered in the green. A. RGB images taken directly from the camera. Color channels are separated. A. Green Channel. The coleoptile is a white organ, it appears green under green light. The markers do not modify the apparent shape of the object (no holes). Under green light, the orange fluorescent marker emits light at the same time in the green and red channel. C. In the red channel, only the orange fluorescent markers are visible.

Prepared coleoptiles were transferred to an experimental darkroom, positioned 80 cm in front of a camera *(Olympus SP-550UZ* or *SP-560UZ)*, and oriented in profile to the camera lens such that maximum curvature was recorded and no part of the coleoptile structure was obscured by the seed. Photographs were taken every 15 minutes during a 24 hours period, with the camera’s flashlight covered by a “safe” green light filter *(Lee 139 Primary Green).* Preliminary experiments showed that neither the pigment markers nor the flash altered any aspects of coleoptile movement and growth.

### Experimental Procedure

Each coleoptile was randomly assigned to one of two treatment groups. In the tilted group, immediately prior to recording, coleoptiles were reoriented from the vertical at a 90° angle (horizontal) and kept in that position for the duration of the experiment. Coleoptiles assigned to the straight (untilted) group were remained vertical, aligned with the direction of tof the gravity field. This second group served as a control to monitor how the aerial organ would grow without gravitropic perturbation.

### Extracting Coleotpile Kinematics

For each coleoptile, the KymoRod software [22] was used to extract movement kinematics from the images recorded during the experiment. From every image taken at time *t*, KymoRod extracted the total length of the coleoptile organ *L*(*t*), and determined each position *s* along the length of the organ from base to apex. For each position s, the organ’s orientation from the vertical *A*(*s, t*) was computed in radians (see Figure 2).

The curvature of the organ *C*(*s, t*) was then calculated with a spatial derivation of the organ’s orientation from vertical as shown:

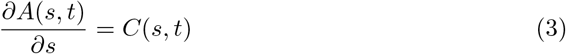

To fully describe the kinematics, the curvature variation 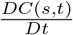 was computed in order to account for organ deformation and elongation [2, 3, 5]. 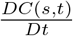 was calculated using the material derivative 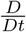, co-moving with each element of the organ derivative. The material derivative of the local variable *C*(*s, t*) is therefore given by:

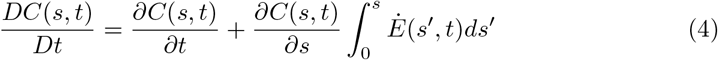

where the sign of 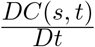 describes the direction of curvature relative to the vertical and *Ė*(*s, t*) is the relative elongation growth rate (REGR) in the direction of the main axis of the organ. *Ė*(*s, t*) was determined using subpixellar displacement mapping function based on digital image correlation [22] which works by comparing target features from sets of digital images. At each position *s* along the length of the organ, we quantified REGR by applying a focal moving window that identified unique patterns of fluorescent markers and assessed marker displacement over time by comparing corresponding windows from sequential images. To yield a single value of elongation growth and curvature variation per coleoptile, we calculated the temporal averages of *Ė*(*s, t*) and 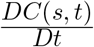 based on the absolute values of the entire set of kinematic extracted from each coleoptile.

### Calculating the Balance Number B

For all coleoptiles in the tilted condition, the balance number *B* was extracted through morphometric measurement as detailed in [14],

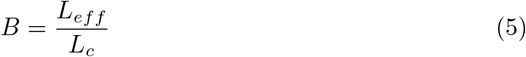

where *B* represents the ratio between the effective length of the growth zone *L_eff_*, and the length of the curved zone *L_c_* at the end of the experiment when no variation of curvature is observed 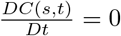. By calculating *B* based on initial morphometrics, we predicted which coleoptiles would ultimately overshoot the vertical. We demonstrated that plants with *B* > 2.8 should display an overshoot during the gravitropic movement [14]. Because *B* is a measure of the gravitropic movement, it cannot be calculated for coleoptiles in the straight and vertical condition.

### Characterizing Oscillations

The relative elongation growth rate *Ė*(*s, t*) and the curvature variation 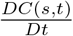 were represented on spatiotemporal kymographs. Thresholding was used to identify the positions of oscillation maximums and minimums. The oscillation period *T_p_* was defined as the time *T* between *n* non-successive peaks (maximums) as shown in:

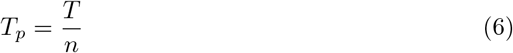

The velocity of these oscillation “pulses” was determined by first fitting a line that followed the position of each oscillation maximum (Figure 5) so that the position of each peak along the organ at time *t* is described by *P* (*t*). Taking into account the time since the pulse start *t_p_*, pulse velocity is given by

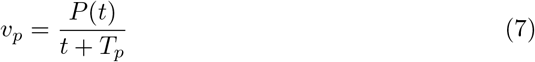

**Fig 5.**
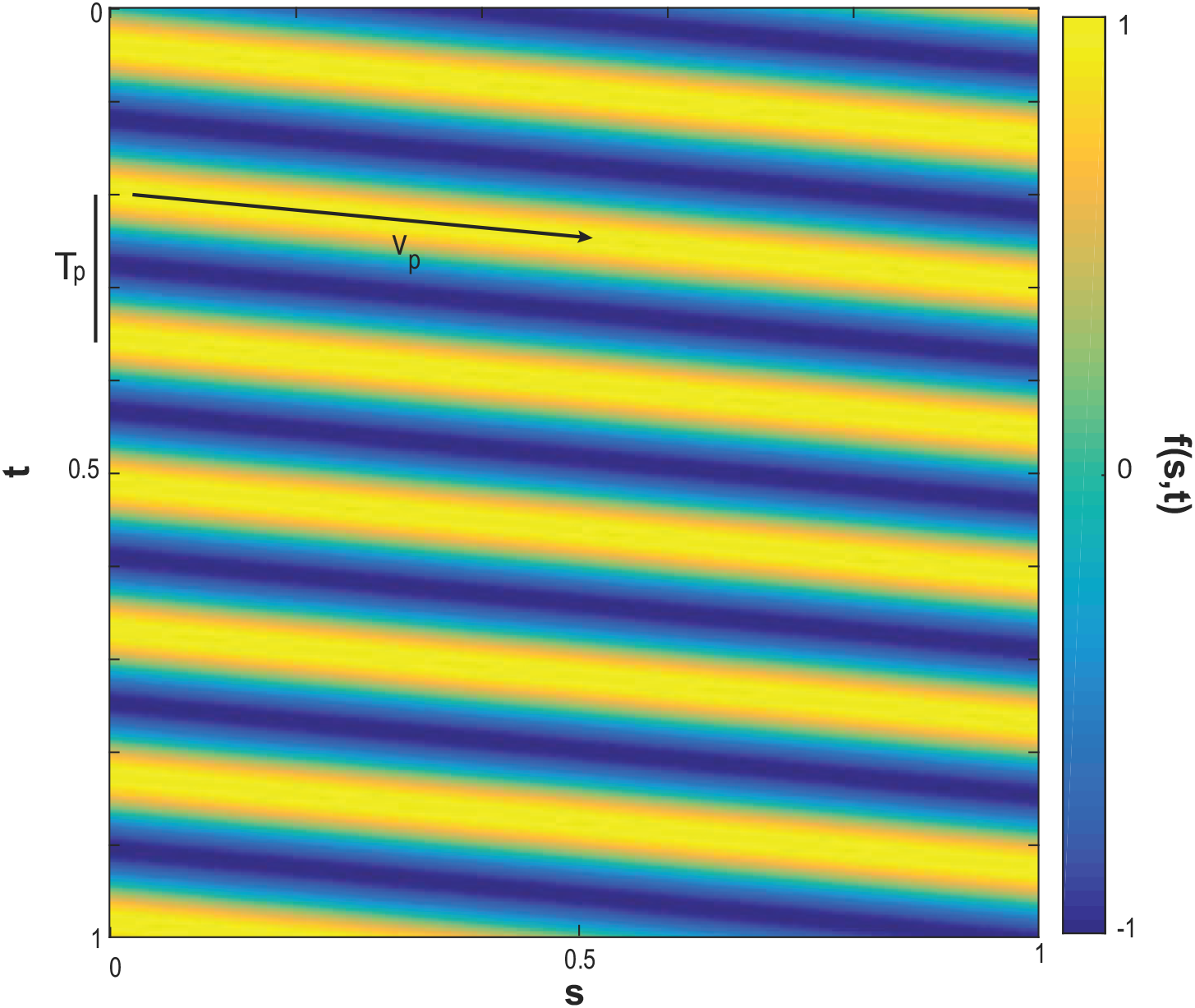
An idealized propagating wave, 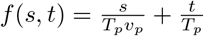. The values of the function *f*(*s, t*) with respect to time *t* and curvilinear abscissa *s*.

### Data Analysis

All statistical and image analyses were conducted in MATLAB (v. 9.0 R2016a, Mathworks Inc.). After calculating the balance number B for each coleoptile in the tilted group, we compared overshoot model predictions [14] to actual experimental data using Fisher’s Exact test to quantify if the model was accurately able to predict coleoptile overshoot. We tested for differences between coleoptiles in the tilted and straight experimental treatments using Mann Whitney U tests and compared: the oscillatory periods, velocity of oscillatory pulses, and temporal averages, of both elongation growth rate *Ė*(*s, t*) and curvature variation 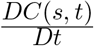. To uncover if oscillations in elongation growth and curvature variation were somehow related, and if this relationship depended on tilting treatment, we calculated the image correlation statistic *r* between these two oscillation kymographs for each coleoptile. We then compared the distributions of r from coleoptiles in the straight and tilted treatments with a Mann Whitney U tests. Movies of all plants analyzed are available online [23, 24].

## Results

Full movement dynamics were extracted from 62 coleoptiles in the straight condition, and 54 coleoptiles in the tilted condition. The unequal sample size is due to several coleoptiles becoming “pierced” by the plant’s initial leaf within the recording time-frame of the experiment (see [8, 9]). As the piercing introduces new movement dynamics to the coleoptile, data from these individuals was discarded.

### Description of movement patterns

The straightening movement of a tilted wheat coleoptile follows the generic pattern of gravitropic movement previously formalized in the AC model [14] (Fig. 3). After being tilted, the coleoptile initially responds by curving the entire aerial organ towards the vertical. Once the tip or apex of the coleoptile has reached the vertical, a process that take anywhere from 1 to 3 hours, the curvature concentrates near the base of the aerial organ while the remainder of the coleoptile continues to elongate along the vertical (Figure 6).

**Fig 6.**
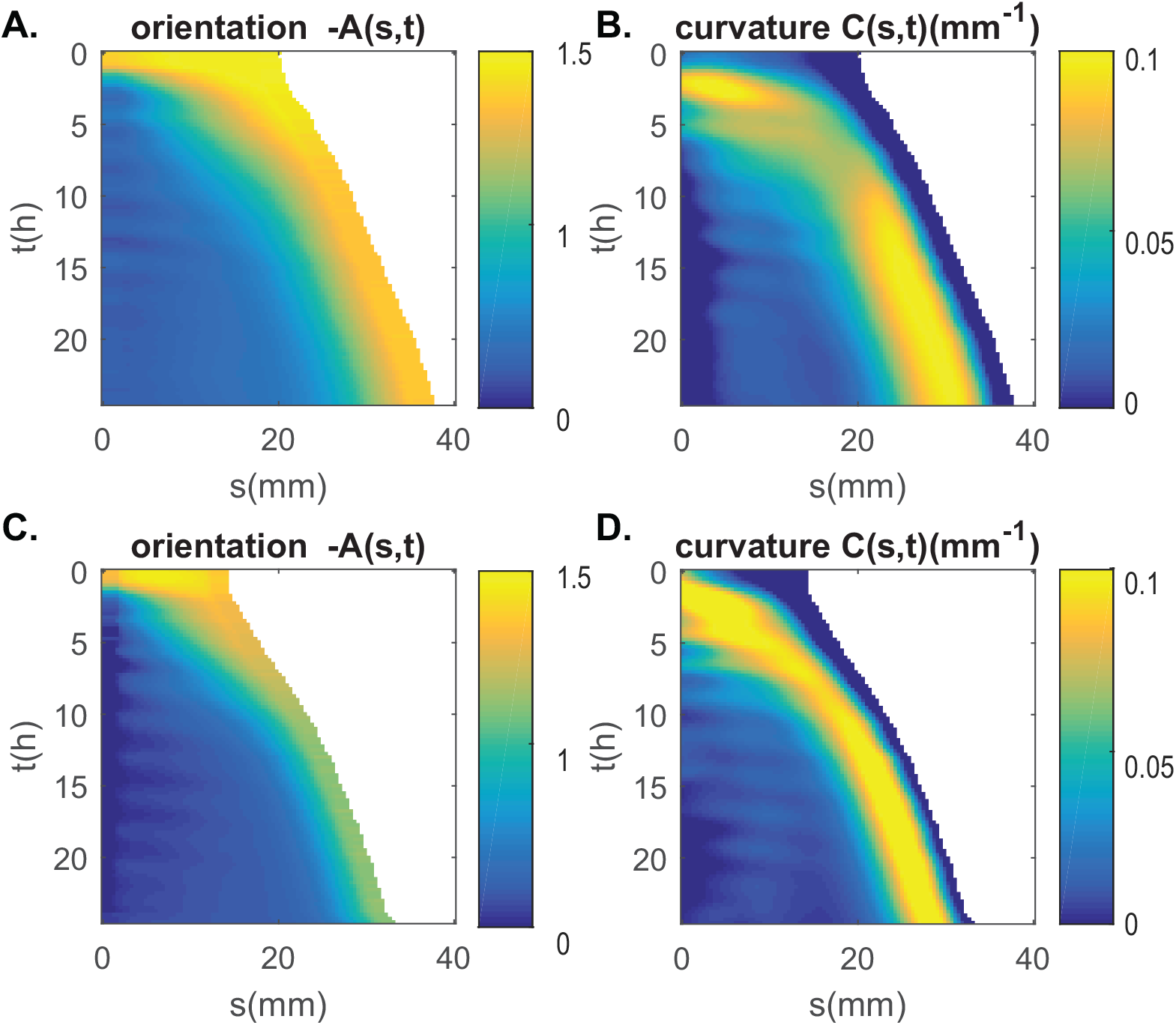
Kinematics of gravitropic movement of two wheat coleoptiles (A-B from Fig. 1) shown as color maps (kymograph) A-C. the orientation *A*(*s, t*) and B-D. the curvature with respect to time *t* and curvilinear abscissa *s* (the arc length along the median measured from the apex *s* = 0 to the base of the organ *s* = *L*). As the coleoptile is elongating, the area covered is increasing with the time. The angle is measured from the vertical in radians. The curvature is measured in *mm*^−1^.

### Growth regulation in wheat coleoptiles exceeds AC model predictions

Fewer tilted coleoptiles overshot the vertical during our experiment (*n_observed_* = 5) than predicted by the AC model (*n_prodicted_* = 16), both during responsive movement and in their final shape (Figure 7). Our Fisher’s Exact test revealed that the difference between predicted and actual overshoots was significant (*n* = 54, *df* = 1, *p* = 0.0069, *OddsRatio* = 4.12, 95% *Cl* [0.99,17.28]), indicating that the tilted coleoptiles in this experiment possess a regulatory mechanism of gravitropic movement that is currently unrepresented in the AC model.

**Fig 7.**
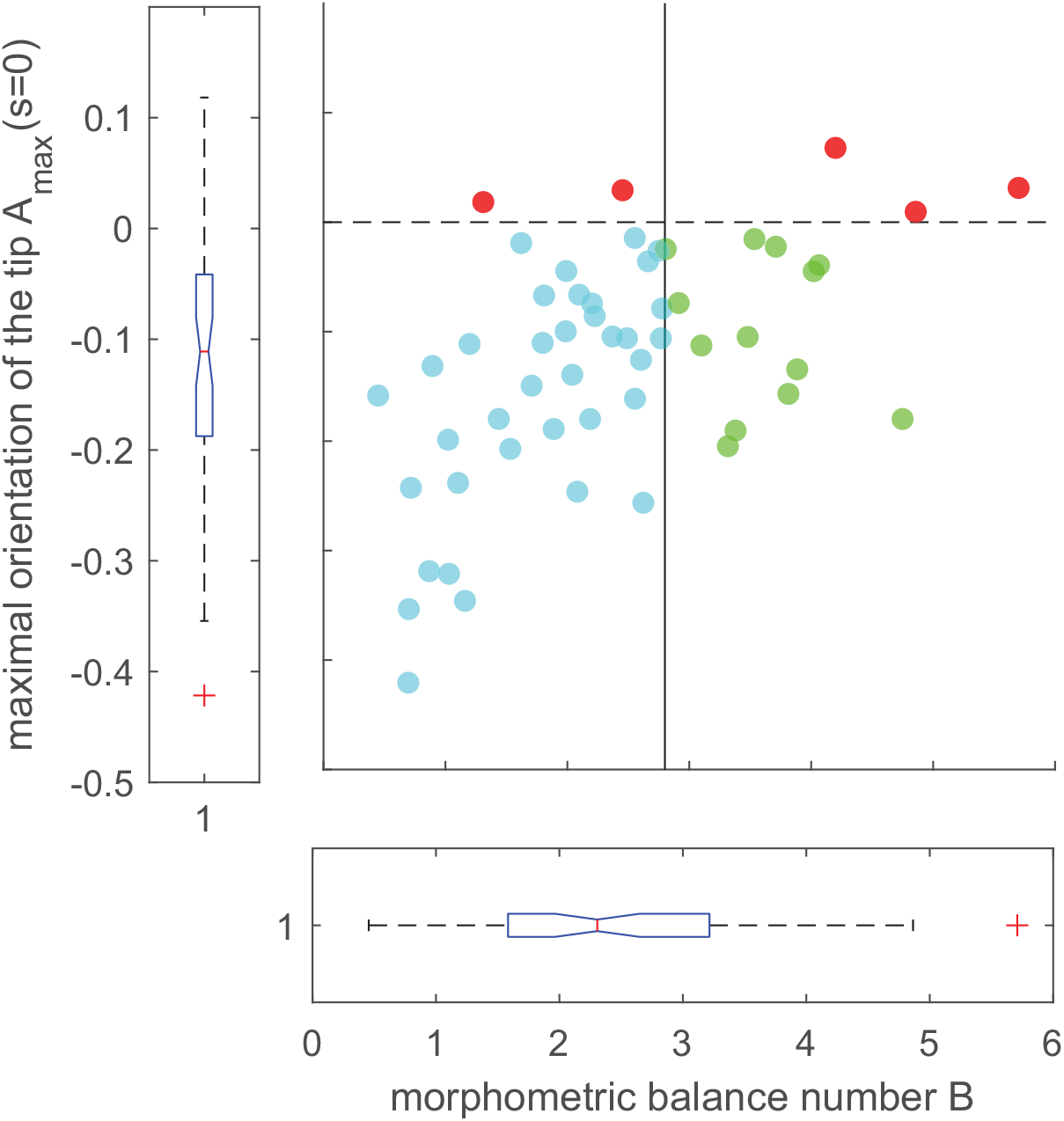
The maximal orientation of the tip, *A_max_*(*s* = 0), as a function of the associated morphometric metric balance number *B* extracted from experiments. The vertical line (*B* = 2.8) shows the transition between the plants that are predicted to not overshoot the vertical (on the left, in blue) and the plants that are predicted to overshoot the vertical (on the right, in green). The dashed shows the transition with the plants that have been observed to overshoot the vertical (on top, in red).

### Whole-plant kinematics: Relative Elongation Growth Rate

The tilting manipulation did affect elongation growth rate with straight coleoptiles having a slightly higher average growth rate than tilted coleoptiles (Med. Straight = 1.0 × 10^−5^*s*^−1^, Med. Tilted = 9.3 × 10^−6^*s*^−1^, *U* = 2896, *p* = 0.019). As evidenced by the kymographs shown in Figure 8, elongation growth rate showed a pattern of oscillation, with growth peaks passing through the aerial organ in waves. However, there was no difference in the average duration (period) of oscillations (Med. Straight = 2.9*h*, Med. Tilted = 2.8*h, U* = 1084, *p* = 0.88) and no difference in the velocity of oscillation pulses along the length of the plant organ (Med. Straight = 11.3*mm.s*^−1^, Med. Tilted = 12.3*mm.s*^−1^, *U* = 2.64 × 10^4^, *p* = 0.12) between the straight and tilted coleoptile groups (Figure 9).

**Fig 8.**
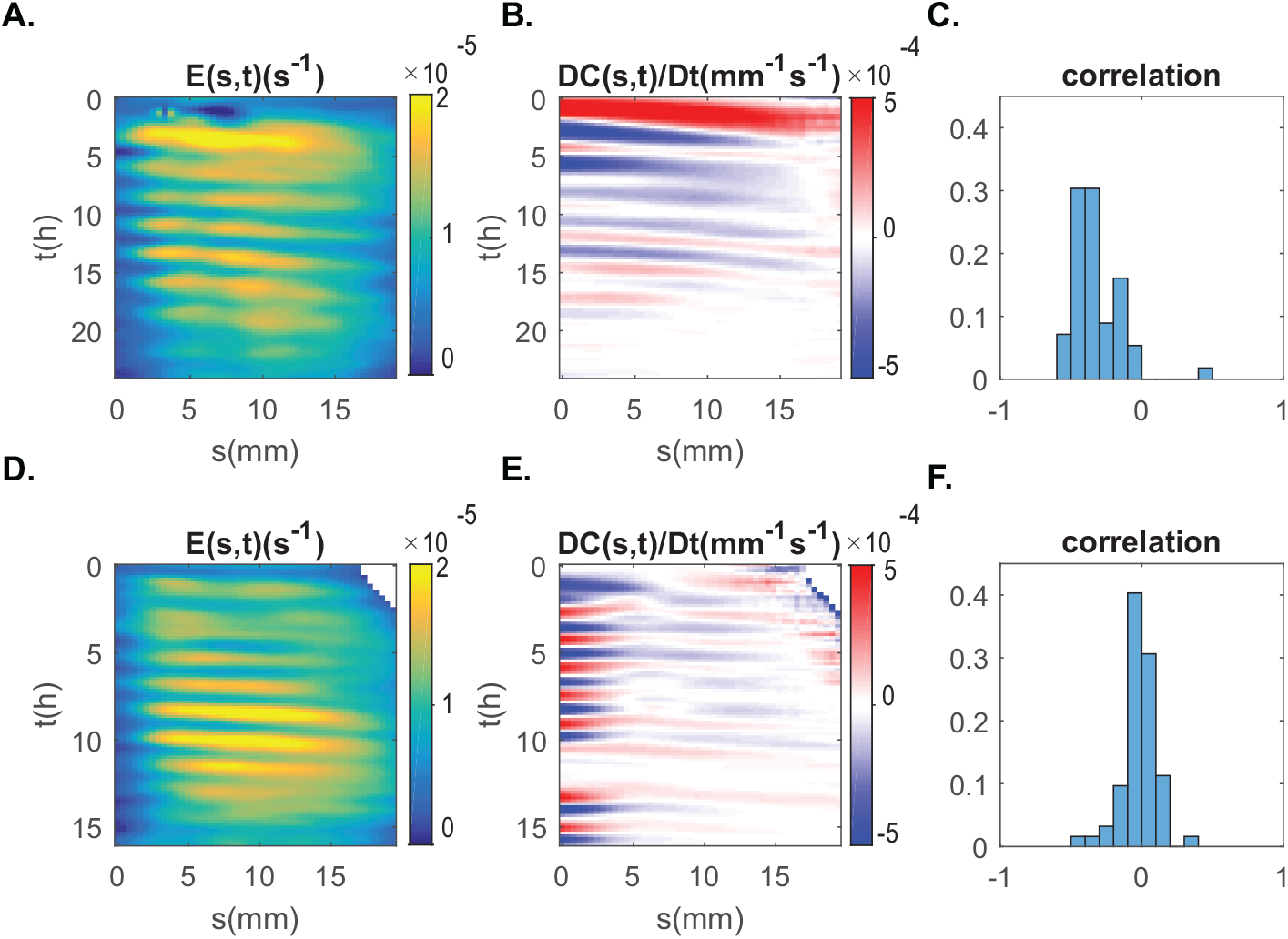
Relative Elongation Growth Rate *Ė*(*s, t*) and curvature variation *DC(s, t)/Dt* of two indivdual coleoptiles. A-B. untilted coleoptile, D-E coleoptile been tilted at 90° from the vertical. Color maps plotting *Ė*(*s, t*) in s^−1^ (A and D) and the curvature variation *DC(s, t)/Dt* in *mm*^−1^ *s*^−1^ (B and C) with respect to time *t* and curvilinear abscissa s (the arc length along the median measured from the apex *s* = 0 to the base of the organ *s* = *L*). Distribution of correlation between *Ė*(*s, t*) and *DC(s, t)/Dt* for all experiments (C. untilted and F. tilted)

**Fig 9.**
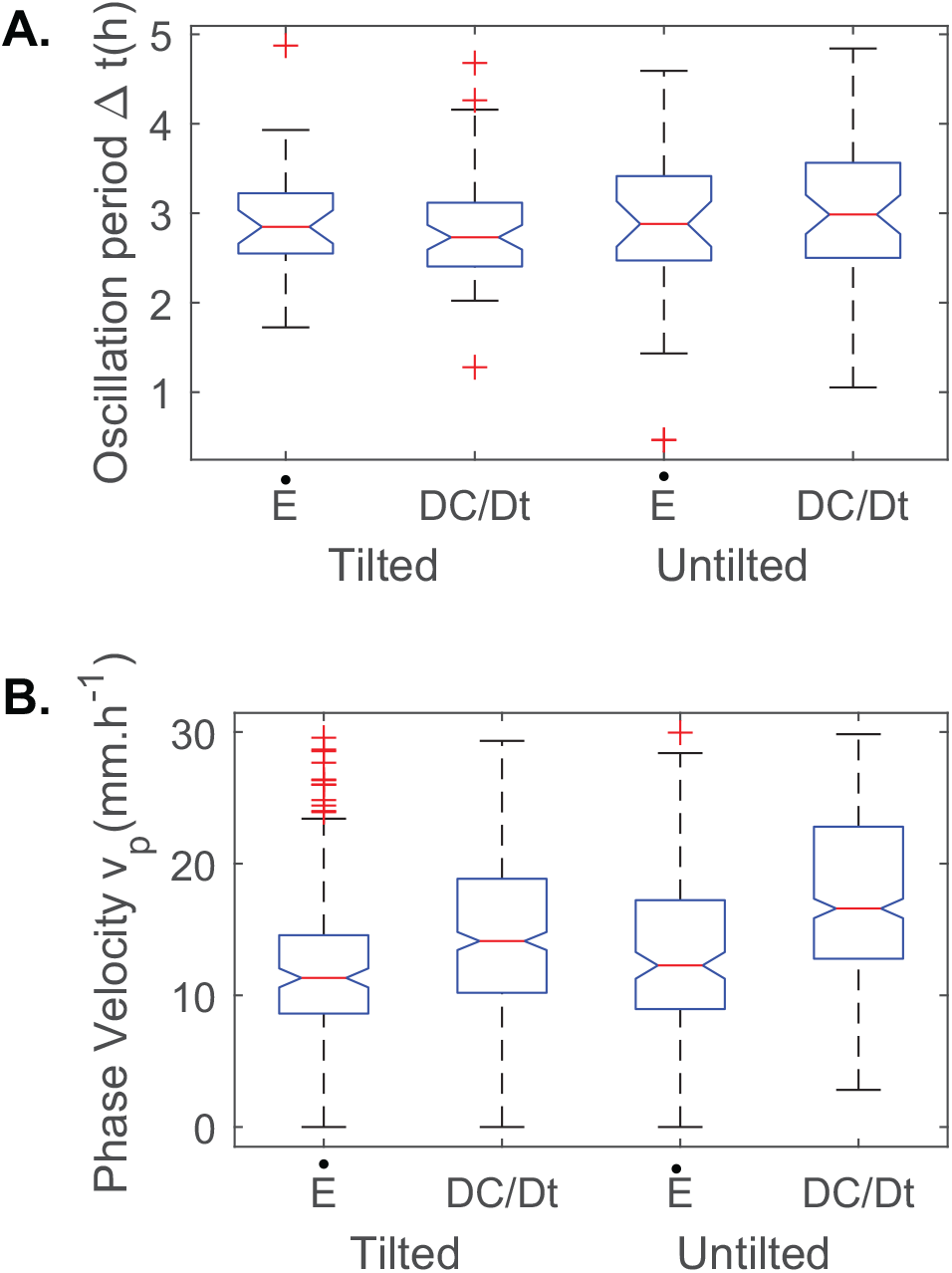
Distribution of the wave propagation parameters. A. Ocilllation period Δ*t*(*h*) in hours and B. Phase velocity *υ_p_* in *mm.h*^−^1 measured for the elongation *Ė* and the curvature variation *DC/Dt* for non-tilted coleoptiles and tilted ones.

### Whole-plant kinematics: Change in curvature

Tilted coleoptiles had a significantly higher average curvature variation than straight plants (Med. Straight = 4.36 × 10^−5^*mm*^−1^*s*^−1^, Med. Tilted = 6.86 × 10^−5^*mm*^−1^ *s*^−1^, *U* = 8331, *p* = 8.74 × 10^−4^). Oscillation patterns were also present in kymographs of curvature variation (Figure 8). Between coleoptiles from the straight and tilted groups there significant differences in oscillation period (Med. Straight = 2.8*h*, Med. Tilted = 3.0*h*, *U* = 3709, *p* = 0.038) and in the velocity of these pulses (Med. Straight = 14.1*mm.s*^−1^, Med. Tilted = 16.6*mm.s*^−1^, *U* = 144624, *p* < 0.001), with tilted coleoptiles demonstrating oscillations that were shorter and travelled more slowly along the aerial organ (Figure 9).

### Tilted coleoptiles show correlated waves of elongation growth and curvature change

While coleoptiles in the straight treatment showed no evidence of a relationship between oscillations in elongation growth rate and curvature variation on average, tilted coleoptiles demonstrated a significant relationship between the two oscillatory manifestations (Med. Straight = —0.026, Med. Tilted = –0.33, *U* = 1845, *p* < 0.001) (Figure 8). This relationship is described by the data as a moderately strong and negative, indicating an asynchronous regulatory mechanism that causes the oscillations of elongation growth and curvature variation to be shifted and slightly out of phase.

## Discussion

The process of gravitropic movement in plant aerial organs was recently formalized in a unifying model called the “AC model” [14]. Yet, while this model often predicts that plants will “overshoot” the vertical, many plants have been observed to demonstrate a greater regulation of gravitropic movement than currently described by this model and remain vertical during growth [5, 14]. To discover the additional movement regulations that prevent plants from overshooting the vertical, we took two groups of coleoptiles and subjected coleoptiles in one of the groups to a gravitropic perturbation (tilted), while leaving coleoptiles in the second group unaffected (untilted) to serve as a control group. Using the kymorod software [22], we extracted the complete movement kinematics of each coleoptile from both groups, in particular focusing on the relative elongation growth rate (REGR) and curvature variation for each coleoptile.

To determine if our sample population possessed this greater regulation described above, we first used these morphometric measures to calculate a “balance number” or B value [14] for each coleoptile from the tilted group. By comparing these B values to previous results from the AC model, we predicted which coleoptiles form the tilted condition would overshoot the vertical. Our data demonstrated that fewer coleoptiles than predicted overshot the vertical (Figure 7), strongly suggesting the presence of additional gravitropic movement regulation in our coleoptile sample.

We then attempted to find this regulatory mechanism by comparing the kinematics of tilted and untilted coleoptiles. First, we examined average measures of elongation growth rate and found that straight coleoptiles had a slightly higher average REGR than those in the tilted group. When looking at a temporal plot of REGR over time (Figure 10), we observed that tilted plants had a slightly lower REGR at the beginning of the experiment, but that this difference quickly dissipated over time. This result is not surprising as tilting a plant disrupts the flow of the growth hormone auxin, causing a temporary reduction in elongation and growth [5]. When we compared average curvature variation between groups, we found a highly significant difference in curvature variation, with tilted coleoptiles displaying much higher values of curvature than controls. Given the nature of our gravitropic treatment, this difference was expected.

**Fig 10.**
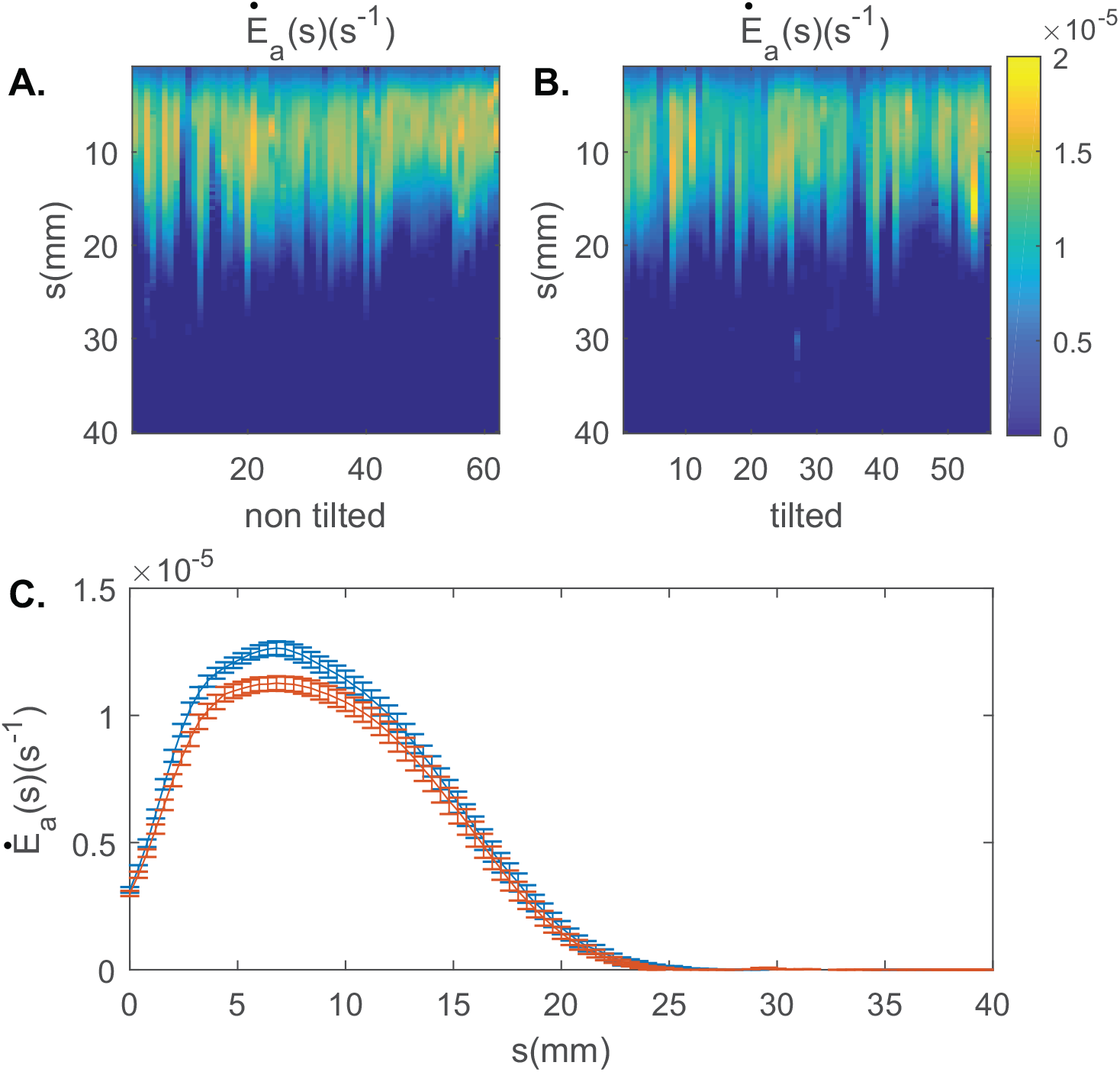
Relative Elongation Growth Rate averaged on the whole experiment for each individual coleotpile A. non tilted coleoptiles and B. tilted coleoptiles C. Relative Elongation Growth Rate averaged for all individuals in red, the coleoptile is not tilted and vertical, in blue the coleoptile is tilted 90° from the vertical.

Closer temporal examination of the morphometric data revealed pulses along the entire length of the aerial organ in elongation growth rate and variation of curvature over time (Figures 6, 8). By tracking the maxima of these pulses over time, we observed that these oscillations in elongation growth rate and variation of curvature travelled from the apex to the base of the coleoptile. Other oscillations in different forms of plant growth have been described in the literature [25], but these reports are infrequent and the features of these oscillations are clearly different from what we have observed in the current study. indeed most of the growth oscillations reported in the literature so far occur during the nutation of the aerial organ, e.g. in sunflowers [26], or in very particular plant organs/cells such as pollen tubes and root hairs [21, 27, 28]. When oscillations of differential growth were observed during plant nutation, the oscillations were identified by comparing growth rates on opposing lateral sides of the plant organ [26]. In plant nutation, unequal lateral distribution of auxin during rapid growth results in successive differential elongation on opposing sides of the aerial organ, and during such “rotation of growth” one would expect to see opposing lateral oscillations. However, our observed oscillations traveled from the apex of the aerial organ to the base along the median line, and so cannot be accounted for by plant nutation. In addition, the oscillations of growth that have been described in pollen tubes and root hairs [27, 28] are limited to occurring at the very tips of these specialized organs. This scale and localization of growth oscillations are completely different from those observed in our coleoptiles, which occurred along the entire length of the aerial organ.

Neither oscillation period, nor oscillation velocity in REGR significantly differed between tilted and untilted coleoptiles, demonstrating that our gravitropic perturbation had no effect on the oscillatory patterns of REGR. A potential candidate to explain the REGR pulses that were observed in both groups of coleoptiles is auxin, as well-known and major growth factor in plants. The average velocity of propagation of auxin in coleoptiles (12*mm.h*^−1^ according to [29, 30] is compatible with our own measurements (*υp* = 11.3 ± 6.0*mm.h*^−1^ for straight coleoptiles and *υp* = 12.3 ± 6.6*mm.h*^−1^ for tilted coleoptiles). However, the average period of time between two pulses in our data set (*tp* = 2.2*h*) is four for five times longer than the pulses of auxin that have been previously recorded in coleoptiles [31–33]. Therefore, while auxin may be driving some aspects of the REGR oscillations in our coleoptiles, the previously known dynamics of auxin can’t completely explain the presence and features of these oscillations.

While gravitropic treatment group did not affect features of REGR oscillations, it did affect features of oscillations in curvature variation. Between coleoptiles from the straight and tilted groups there were significant differences in oscillation period and the velocity of oscillation pulses, with tilted coleoptiles demonstrating oscillations with shorter periods that travelled more slowly along the aerial organ. Similar oscillations of curvature variation have been previously identified during gravitropic movement [19]. But no conclusive mechanisms were found to explain these oscillations as the study relied only on apical tracking, a method of gathering global measurements that can neglect more detailed and local movement dynamics (see [21] for a discussion on the limits of apical tracking).

Most interestingly, we observed a relationship between oscillations in REGR and curvature variation in tilted coleoptiles (Figure 8), suggesting that tilted coleoptiles had greater couplingn between curvature and REGR than straight coleoptiles. This relationship was moderately strong and negative, demonstrating that regions of lower REGR were associated with higher variation of curvature, and indicating an asynchronous regulatory mechanism that causes the oscillations of elongation growth and curvature variation to be shifted and slightly out of phase. Local curvature and growth are indeed known to be correlated in plants, and curvature variation can be expressed as a function of the elongation rate:

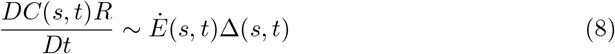

where Δ(*s, t*) accounts for the distribution of the differential growth on each side of the coleoptile during elongation [5]. As no global shrinking of plant aerial organs are expected during growth, elongation *Ė*(*s, t*) will always have a positive value. Therefore, the local elongation rate can only modulate the amplitude of curvature rate, but not its sign, so elongation alone cannot affect the direction in which the organ is curved. However, we observed both a negative correlation between REGR and curvature variation oscillations and a fewer than predicted number of overshoots. Combined, these two results indicate the presence of an additional mechanism that is capable of affecting the relationship between local growth and curvature, as well as the sign of curvature rate. As the differential of growth Δ(s, t) is the remaining term affecting elongation rate, this strongly suggests the presence of a term and/or mechanism influencing the correlation between elongation rate and the distribution of differential growth.

A better understanding and more formal description of this mechanism term (currently missing from the AC model), could come from comparing the oscillatory behaviors of tilted coleoptiles that did and did not overshoot the vertical. Indeed, greater experimental examination would let us know if and how this regulation changes with variation in non-verticality. However, in the current study the actual number of overshoots was incredibly small, and did not provide us with a sample size large enough for statistical comparison with tilted coleoptiles that failed to overshoot the vertical. Given the low rate of overshoots, acquiring a sufficient sample size would require hundreds, perhaps thousands, of wheat coleoptiles to be tested. This approach could also allow us to determine if correlated oscillations in REGR and curvature variation are a general phenomenon among plant species.

In order to explore the REGR variations coupling mechanisms involved in gravitropic regulation, the number of overshooting cases should be increased. One may explore the natural genetic variability in that purpose. Additionnally mutants may be considered. In the wheat background the available mutants are in a small number and a a specific screening should be achieved. The alternative is to turn to the “model species” of the genomics, such as Arabidopsis. Actin and Myosin mutants demonstrating large and recurrent overshoot have been described by [34], and found to be consistent with a lack in proprioception. But just like in wheat, the mutants should now be screened for more subtle differences in overshoot. In any cases, this work has properly defined the way to proceed and it should be, methodologically speaking, straightforward. This should give the way to further out our understanding of the mechanisms that interact in allowing the plant to control their posture efficiently all along their life.

## Acknowledgments

RB wants to thanks Stéphane Ploquin for his help with the experimental setup. All original data are available at [35]. A series of 4 movies are available. They show the timelapse movement of wheat coleoptiles [8, 9] as well as the kinematics of growth and curvature variation [23, 24].

